# Ecological and evolutionary consequences of selective interspecific information use

**DOI:** 10.1101/2022.03.25.485764

**Authors:** Reetta Hämäläinen, Mira H. Kajanus, Jukka T. Forsman, Sami M. Kivelä, Janne-Tuomas Seppänen, Olli J. Loukola

**Author notes:** Shared first authorship (Reetta Hämäläinen, Mira H. Kajanus). Corresponding authors: Reetta Hämäläinen, PL 8000 FI-90014 Oulun yliopisto, Finland; and Mira Kajanus, PL 8000 FI-90014 Oulun yliopisto, Finland.

## Abstract

The ecology of social information use has been studied in many intra- and interspecific contexts, while the evolutionary consequences of social information use remain less understood. Furthermore, *selective* social information use, where individuals are discriminative in their decision-making on how to use social information, has been overlooked in interspecific context. In particular, the intentional decision to reject a behavioural trait observed via social information, has gained less attention, although it has recently been shown to occur in various taxa. We develop an individual-based simulation model to explore in which circumstances social information use leads to different coevolutionary outcomes among populations of two species. The initial phenotypes and the balance between costs of competition and benefits of social information use determine whether selection leads to trait divergence, convergence or coevolutionary arms race between two species. Based on existing literature, we propose that selective decisions of individuals, including active rejection, may have far-reaching fitness consequences, potentially leading to similar evolutionary consequences among the populations of the information source and the user as predicted by our model. Overall, we argue that the eco-evolutionary consequences of selective interspecific social information use may be much more prevalent than thus far considered.

## 1. Introduction

Animals need to make decisions throughout their life cycle, and the decisions (e.g., about forage and breeding sites or breeding investment) often have fitness consequences. To make responsive decisions, animals need information about their ecological or social environment. Information that individuals obtain may include directly acquired personal information, information transferred through parental effects or information acquired in social context from con- or heterospecifics (Danchin *et al*. 2004; Seppänen *et al*. 2007; Farine *et al*. 2015). Acquisition of **social information** (see Glossary) may be faster and less costly than gathering information merely via own observations (Dall *et al*. 2005). Study of social information use among conspecifics has already a long tradition in ecology and behaviour (Kawamura 1959; Galef 1995; e.g., Danchin *et al*. 2004; Seppänen *et al*. 2007), yet the overall importance of including social information in ecological theories, such as competition theory (Gil *et al*. 2019) and community ecology (O’Connor *et al*. 2019), has only recently gained more attention.

The concept of social information use consists of the source of the information (hereafter **information source**), social information that the information source produces and a user of that information (hereafter **information user**). Social information is observable for con- and heterospecifics, examples including signs of parasitization on other individuals (Loukola *et al*. 2020b) and outcomes of reproductive decisions (e.g., location of a nesting site [Forsman et al. 2002; Schmidt et al. 2015; Chiatante 2019]; and reproductive success [Forsman & Seppänen 2011; Seppänen et al. 2011; Schuett et al. 2017]). Individuals may switch between social and directly acquired personal information adaptively rather than use a fixed strategy. For example, in situations where the individual is older, more experienced or has different phenology than the source, it should ignore the social information, and instead rely on innate responses, past experiences and personal information (Galef & Laland 2005; Schmidt *et al*. 2015; Kendal *et al*. 2018; Laland *et al*. 2020). Additionally, individuals may need to be selective in the way they use social information. Traditionally, studies on social information use have concentrated merely on the selective copying-decisions (e.g., when or who to copy) of the information user (Laland 2004; Enquist *et al*. 2007). However, individuals can choose to copy the **observed behaviour** or actively reject it (see Box 1 and Fig. 1 for more detailed description). This active rejection of observed behaviour must not be confused with resource partitioning, which may also be mediated by social information use (e.g., Templeton *et al*. 2017). Active rejection may be challenging to distinguish from ignoring an observed behaviour and requires a specific experimental set-up to be identified. Experimental studies where the decision is forced between only two different choices based on the observed decision and the outcome of that decision of another individual, for example a choice between two nest-site characteristics (decision) and clutch size (outcome; Loukola *et al*. 2012, 2013, 2020b) have been used to demonstrate rejection behaviour. When the decision made by the information source is rejected, the opposite behaviour occurs more often than would be expected by random. Selective social information use may increase the fitness of the information user when merely copying the behaviours of others is not always optimal (Laland 2004; Enquist *et al*. 2007).

### Box 1. Active rejection

An individual may selectively copy or reject an observed behaviour (e.g., choice of prey) of a con- or heterospecific based on the outcome of this behaviour (e.g., breeding success; Fig. 1). If the observed individual (information source) has made a poor decision leading to seemingly negative outcome, rejecting this decision should be more advantageous than copying it (Forsman *et al*. 2018). Seppänen et al. (2007) states: “Active rejection of behaviours of poor individuals can facilitate decision making by reducing the set of alternatives to choose from, thus reducing uncertainty. Especially when the number of alternatives to choose from is small, being able to discard even one of the alternatives provides considerable advantage. In the case of just two alternatives (a binary choice), rejection of the alternative exhibited by a poor individual leaves just one alternative to be adopted”. Accordingly, several binary choice experiments in birds (Seppänen *et al*. 2011; Loukola *et al*. 2013; Hämäläinen *et al*. 2021a) and bees (Loukola *et al*. 2020b; Romero-González *et al*. 2020a) have shown active rejection rather than ignoring the observed behaviour.

A prerequisite for SIIU and rejection is the reliability of the information source (Loukola *et al*. 2013; Forsman *et al*. 2018). In birds, individuals with good problem-solving skills have larger clutch sizes than individuals with lower problem-solving skills regardless of habitat quality (Cole *et al*. 2012). If breeding success can be a reliable cue of the cognitive abilities of the information source, then high and low fitness of individuals should reflect good and poor decision-making abilities, respectively. Hence, a successful individual should be copied (Loukola *et al*. 2013, 2020b; Romero-González *et al*. 2020a), while it would be better to reject the decisions demonstrated by unsuccessful individuals (Loukola *et al*. 2013, 2020b). Bumblebees (*Bombus terrestris*) are selective in information use in relation to learned reliability of information source; they prefer to land on flowers, that are near the individuals whose presence had previously predicted reward, indicating copying the behaviour of seemingly successful individuals (Romero-González *et al*. 2020a).

The concept of loss aversion could be a possible rationale behind rejection behaviour. Loss aversion predicts that there is a tendency to prefer avoiding losses to acquiring equivalent gains (Tversky & Kahneman 1992). In animals, the relevant currency, is fitness or reliable fitness correlate. Copying an unreliable or unsuccessful information source would risk low fitness, so, rejecting choices of unsuccessful individuals reduces the risk of low fitness. In passerine birds, rejecting the apparent choices of unsuccessful information source birds that plausibly make poor decisions is stronger than copying the choices of successful information sources that apparently have made good decisions (Forsman & Seppänen 2011; Seppänen *et al*. 2011; Loukola *et al*. 2013). In addition, it has been proposed that information acquisition can be seen as a type of evolutionary bet hedging and avoiding poor choices is more important for the evolution of information acquisition strategies than making the very best choices (Forsman & Kivelä 2021).

The concept of rejection is an integral part of SIIU and, together with copying, it facilitates decision-making in opposite directions, depending on the perceived fitness-consequences of the alternative choices. Still, rejection has not been included in the current theory of social information use, although it has been recorded in individual studies among conspecifics (Kendal *et al*. 2009; Kurvers *et al*. 2010; Loukola *et al*. 2012; Loyau *et al*. 2012; Szymkowiak *et al*. 2016) and heterospecifics (Seppänen *et al*. 2011; Loukola *et al*. 2013, 2020b; Tolvanen *et al*. 2018; Morinay *et al*. 2020; Hämäläinen *et al*. 2021a).

**Figure 1.**
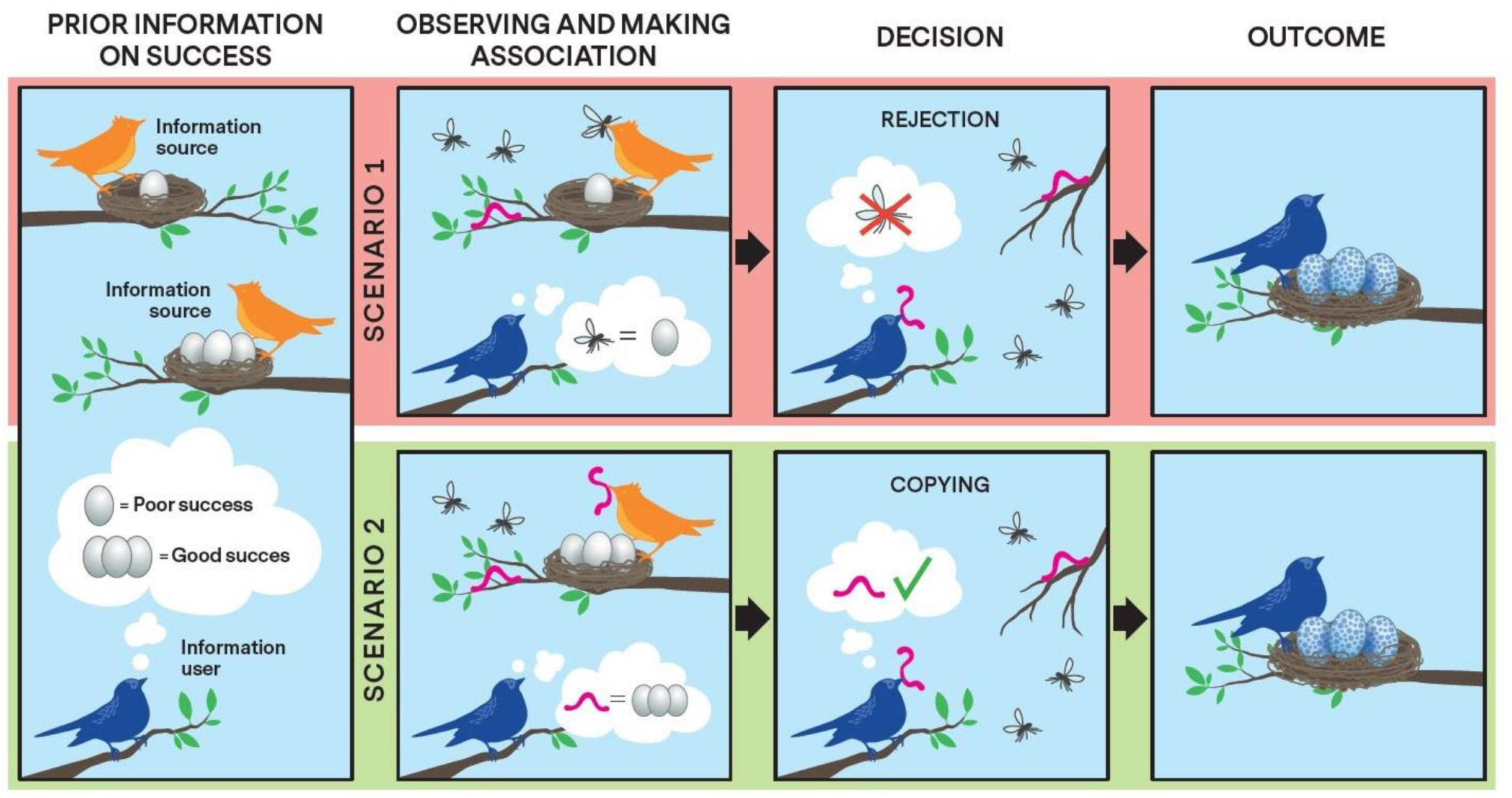
There may not be a direct cause-effect chain of reasoning in selective social information use, but the information user may affiliate all the observed behaviours of the information source to the success of the source. Figure 1 is a schematic illustration showing the process of selective information use for discriminatively deciding to copy or reject a single observed behaviour, using breeding success as the cue from the information source (orange bird) and prey choice as the decision of the information user (blue bird). The information user gains the information of poor- or high-quality information source (con- or heterospecific) from a cue, such as breeding success (clutch size, one or three eggs; PRIOR INFORMATION ON SUCCESS). In **Scenario 1** (top row), the information user observes the information source with low breeding success and associates the observed prey choice (mosquito) of the information source with low breeding success of the source (OBSERVING AND MAKING ASSOCIATION). The information user then encounters a situation with the same available food resources, and when having to make a decision on which resource to consume, the information user follows the previous association between the prey choice (mosquito) and low breeding success of the information source. Thus, the information user decides to reject (DECISION: REJECTION) the observed behaviour and decides to consume the other available food resource (caterpillar), for which the information user does not have prior information on. This decision likely results in high breeding success of the information user (OUTCOME). In **Scenario 2** (on the bottom row), the information user associates the prey choice (caterpillar) of the information source with high breeding success (OBSERVING AND MAKING ASSOCIATION), the information user decides to copy (DECISION: COPYING) the behaviour of the information source and consumes the same food resource (caterpillar) instead of the food resource without prior information on (mosquito). Also this decision likely results in high breeding success of the information user (OUTCOME).

It is important for the information user to associate the success of the information source to the behaviour of the source to either copy or reject the observed behaviour (Forsman *et al*. 2018; Box 1 and Fig. 1). Information on high fitness (e.g., large clutch size in birds; Seppänen *et al*. 2011; Morinay *et al*. 2020) or successful nesting (e.g., healthy nest in solitary bees; Loukola *et al*. 2020b) of the information source increases the probability that the information user copies the behaviour of the information source, whereas an association with information on low fitness (e.g., small clutch size; Seppänen *et al*. 2011; Loukola *et al*. 2013) or unsuccessful nesting (e.g., parasitized nest in solitary bees; Loukola *et al*. 2020b) increases the probability to reject the behaviour of the information source. Current theory of social information use does not acknowledge the rejection behaviour, even though many case studies show that a selective decision to either copy or reject a certain trait or behaviour could have adaptive consequences (e.g, Loukola *et al*. 2013, 2020b; Forsman *et al*. 2014; Szymkowiak *et al*. 2016; Romero-González *et al*. 2020a; Hämäläinen *et al*. 2021a).

Moreover, there is a well-constructed theory for the adaptive value of *intraspecific* social information use (Laland 2004; Enquist *et al*. 2007; Kendal *et al*. 2009; e.g., Bocedi *et al*. 2012; Loyau *et al*. 2012; Whitehead *et al*. 2019), and despite of the accumulating number of studies over wide range of species (Seppänen *et al*. 2007; Jaakkonen *et al*. 2015; Forsman *et al*. 2018; Whitehead *et al*. 2019; Keen *et al*. 2020; Loukola *et al*. 2020b; Romero-González *et al*. 2020a; Hämäläinen *et al*. 2021a, b), there are gaps in our understanding on the adaptive value of *interspecific* social information use. The fitness consequences of social information use depend on the accuracy of the acquired information and the accuracy decreases with ecological and temporal distance between the information source and information user (Seppänen *et al*. 2007). It has been suggested (Laland 2004) that the adaptiveness of social information use is more likely to result from selective social information use, and we suggest that the adaptive value may be similar across heterospecific as among conspecific information use.

**Selective interspecific information use** (SIIU) has gained increased attention and has been observed in birds (Forsman & Seppänen 2011; Seppänen *et al*. 2011; Loukola *et al*. 2013; Szymkowiak *et al*. 2016; Thorogood *et al*. 2018; Tolvanen *et al*. 2018; Morinay *et al*. 2020; Hämäläinen *et al*. 2021a) and insects (Loukola *et al*. 2020b). This interspecific “reject-the-unsuccessful” strategy (Forsman et al. 2018; Loukola et al. 2020b; Romero-González et al. 2020a; but see Slagsvold & Wiebe 2017) complements existing literature on selective social information use including “copy-the-successful” strategy (Laland 2004). In this study, we focus on copying and rejecting an observed behaviour due to their relevance in ecological and evolutionary consequences of SIIU.

SIIU requires a range of cognitive abilities, which may at first seem beyond those possessed by most animals, but seemingly complex behaviours can be achieved with small brain volumes and explained through associative learning mechanisms (Leadbeater & Chittka 2007; Giurfa 2012; Alem *et al*. 2016; Leadbeater & Dawson 2017; Gatto *et al*. 2021). Insects can display a range of sophisticated behaviours with a miniature sized brain (Leadbeater & Chittka 2007; Avargués-Weber *et al*. 2011; Giurfa 2012; Leadbeater & Dawson 2017; Gatto *et al*. 2021) including concept learning (Giurfa *et al*. 2001), numerical skills like addition and subtraction (Howard *et al*. 2019) and even tool use (Loukola *et al*. 2017). Because insects, that comprise most species in the animal kingdom, show this level of deductive abilities, it is likely that many species through the animal kingdom would be capable for SIIU.

It is known that interspecific social information use affects important aspects of species’ ecology, such as adapting foraging (Farine *et al*. 2015; Keen *et al*. 2020; Romero-González *et al*. 2020b; Hämäläinen *et al*. 2021a), breeding site selection (Kivelä *et al*. 2014; Szymkowiak *et al*. 2017; Chiatante 2019) and dispersal decisions (Bocedi *et al*. 2012; Cayuela *et al*. 2018), leading to population-level consequences (e.g., variation in species’ densities and intensity of competition). These ecological consequences of interspecific social information use may, in turn, affect the fitness effects of social information use and can change the direction (Whitehead *et al*. 2019; Laland *et al*. 2020) or enhance the strength (Danchin *et al*. 2004; Paenke *et al*. 2007; McPeek 2017; Whitehead *et al*. 2019; Laland *et al*. 2020; Martin *et al*. 2021) of natural selection, and potentially affect the differences among species’ traits, highlighting the inherent eco-evolutionary feedback loop in interspecific social information use. Thus, social environment and **trait space** (i.e., multidimensional space determined by both morphological and behavioural traits) changes may be intertwined more than previously considered (Paenke *et al*. 2007; Loukola *et al*. 2013; Magrath *et al*. 2015; Bailey *et al*. 2018; Whitehead *et al*. 2019; Laland *et al*. 2020; Hämäläinen *et al*. 2021b). Still, the effect in evolutionary rates and the full extent of the potential evolutionary consequences of social information use remain poorly understood.

Both the traditional (Brown & Wilson 1956; Hardin 1960; Macarthur & Levins 1967; Pianka 1974) and more recent (Chesson 2000; Costa-Pereira *et al*. 2019; Derbridge & Koprowski 2019; Martin *et al*. 2021; Pastore *et al*. 2021) work on species coexistence expect that mechanisms, such as competition (Pianka 1974; Pastore *et al*. 2021), limiting similarity (Macarthur & Levins 1967; Chesson 2000) or character displacement (Brown & Wilson 1956; Derbridge & Koprowski 2019), lead to **trait divergence** among ecologically similar species. Whether SIIU leads to similar evolutionary consequences remains poorly understood but the short-term fitness benefits of social information use are well-established (Laland 2004; Dall *et al*. 2005; Enquist *et al*. 2007; Kendal *et al*. 2009; Tobias & Seddon 2009; Whitehead *et al*. 2019; Keen *et al*. 2020; Laland *et al*. 2020; Hämäläinen *et al*. 2021b).

On the other hand, through social information use, species with similar resource needs, one or both of the species could benefit from copying the behaviour of the potential competitor, leading to **trait convergence** between the species (Forsman *et al*. 2002; Tobias & Seddon 2009; Seppänen *et al*. 2011; Loukola *et al*. 2013; Tobias *et al*. 2014; Farine *et al*. 2015; Gil *et al*. 2018; Chiatante 2019; Keen *et al*. 2020; Hämäläinen *et al*. 2021b). However, increasing density of con- and heterospecifics plausibly increase competition (Seppänen *et al*. 2007; Forsman *et al*. 2014), and may eventually lead to trait divergence supporting the traditional views of coexistence of similar species (Brown & Wilson 1956; Hardin 1960; Macarthur & Levins 1967; Pianka 1974). Assuming that traits affect resource use overlap between individuals of two species, the difference between the trait values of the information source and the user should have a major effect on the net benefits of SIIU. The balance of costs and benefits can be expected to influence the evolution of the propensity to use social information as well as the direction of trait evolution in the two interacting populations (Lee *et al*. 2016).

We argue that SIIU may result in changes in species’ trait values. Changes in the differences among species’ trait values affect interspecific interactions and so coevolution between species connected by social information use. Thus, divergent resource use decisions could eventually lead to evolutionary trait divergence and convergent resource use decisions to evolutionary trait convergence (Fig. 2). If SIIU and consequent trait convergence result in increasing fitness costs for the information source (i.e., “information parasitism”), then it may eventually lead to evolutionary trait divergence or coevolutionary arms race between the two species. We suggest that SIIU is more common than previously thought and may have major ecological and evolutionary consequences in trait divergence, convergence and arms race between coexisting species.

**Figure 2.**
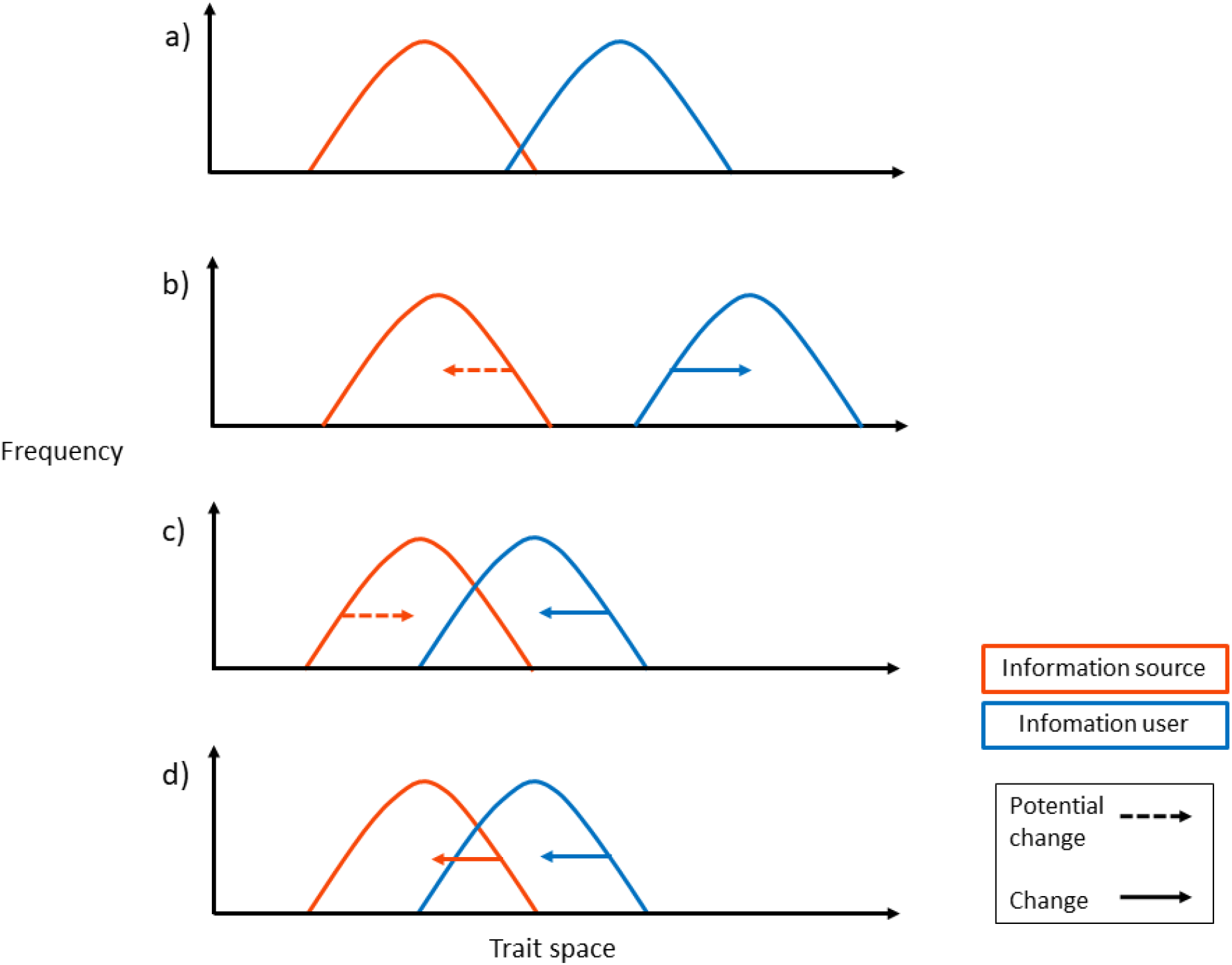
Schematic illustration (adapted from character displacement theory; Brown & Wilson 1956) of the potential population level consequences of selective interspecific information use (SIIU) and competition with the frequency distributions of two coexisting species (the information source in red and the information user in blue) and their hypothetical trait space locations affecting resource use before (a) and after (b-d) SIIU. SIIU may lead to increased competition, where eventually a high level of competition would drive either one or both of the populations to diverge away from each other on the same trait space (b). SIIU may also lead to niche convergence (c), where the information source remains stable on its trait space (or converges towards the information user) and the information user converges towards the information source on the same trait space. Coevolutionary arms race (d) between the two species may occur when the information user evolves towards the trait space of the information source, the information source in turn starts to diverge away from the user to escape from the same trait space (i.e., avoid information parasitism).

The significance of SIIU in determining the location of a species in trait space may be severely underappreciated in the current competition-dominated paradigm. To rigorously explore the potential eco-evolutionary dynamics produced by interspecific social information use and competition, we developed an individual-based simulation model where the initial phenotypes of two species and the balance between the costs of competition and the benefits of social information use affect phenotypic evolution and population dynamics. We also review the importance of the copy/reject behaviour based on empirical studies in existing literature and discuss how these mechanisms may result in the predicted evolutionary outcomes. Finally, we suggest future avenues to address the ecological and evolutionary significance of selective interspecific social information use.

## 2. Modelling eco-evolutionary consequences of interspecific social information use

We developed an individual-based simulation model between interacting populations of two species, A and B, to predict the eco-evolutionary consequences of interspecific social information use. However, to keep the model simple without loss of generality, we did not explicitly include the exact selective decision of the information user (copy/reject) in the model. The distance between the initial phenotypes of the two species and the balance between the costs of competition and the benefits of social information use determine the direction and strength of natural selection. Although the model predictions are derived for social information use, other direct or indirect mechanisms of positive interspecific interactions (e.g., reduced predation pressure) could result in similar evolutionary consequences. In the model, both species reproduce asexually as well as have a semelparous life history and non-overlapping generations. Potential fecundity of individuals belonging to species *i* (*i* = {*A, B*}) is *F_i_*. We consider a phenotype of an individual to represent the trait value (in arbitrary phenotypic units) of any trait (e.g., behaviour, morphology) that may be affected by both competition and social information use.

Realized fecundity of an individual depends on the sizes of the populations of the two interacting species, *N_A_* and *N_B_*. The interspecific component of density dependence is determined by the balance between the costs of competition and the benefits of social information use, both of which depend on the difference between the phenotype of the focal individual and the mean phenotype of the other species. This determination of the phenotypic difference means that each individual of species *i* interacts randomly with the individuals of species *j* (*j* ≠ *i*). We model both the costs of competition and the benefits of social information use with Gaussian functions in relation to the interspecific phenotypic difference. Consequently, density-dependent net effect of interspecific interactions, *α_ki_*, for individual *k*(*k*=1, …, *N_i_*(*t*); *N_i_*(*t*) is the population size of species *i* in generation *t*) of species *i* is

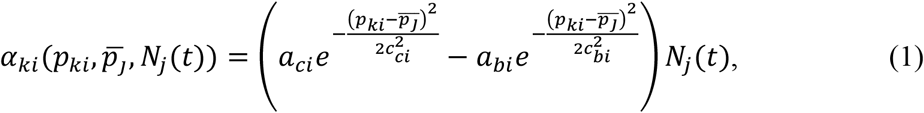

where *p_ki_* is the phenotype (i.e., trait value) of individual *k* of species *i*, 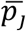 is the mean phenotype of species *j*, and *N_j_*(*t*) is the population size of species *j* in generation *t*. The peak cost of competition, occurring when 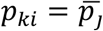, is *a_cj_*, whereas the breadth of the cost function of species *i* in relation to the interspecific phenotypic difference is set by *c_ci_*. Similarly for the benefits of interspecific social information use, *a_bj_* is the maximum benefit at 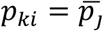, and *c_bi_* determines the breadth of the benefit function. Setting *a_bi_* < *a_ci_* and *c_bi_* > *c_ci_*, there is a net cost to species *i* when interspecific phenotypic difference is small, which changes to a net benefit when the interspecific phenotypic difference increases (see Fig. 3). If *a_bi_* = 0, there is only a cost of interspecific competition that decreases with increasing interspecific phenotypic difference. We assume that the density-dependent costs of intraspecific competition are independent of phenotype (i.e., the focal trait value here) because individuals of the same species will be very close to each other in the trait space even if there would be major variation along one dimension of the trait space. From the above assumptions, the realized fecundity of individual *k* of species *i* in generation *t*, *F’_kit_*, is given by

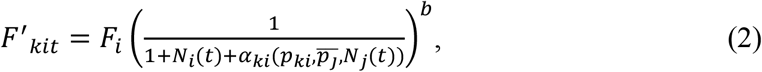

where the exponent *b* determines the strength of joint (i.e., intra- and interspecific) density dependency. Joint density dependency becomes weaker with decreasing *b* (provided that *b* > 0). A net benefit of interspecific social information use results in a negative 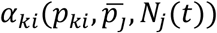, which is why we constrained the sum 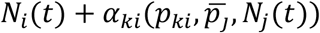 to be ≥ 0, that is, the maximum benefit attainable from interspecific social information use results in the realization of the full reproductive potential. *F’_kit_* was rounded to the closest integer because our individual-based approach required offspring production to consist of full individuals. Population size of species *i* in generation *t* + 1, *N_i_*(*t* + 1), was then calculated as

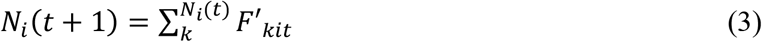

Offspring inherit the phenotype of the mother, but a mutation may take place with a probability of *μ*. Mutation causes a random and normally distributed (~N(0,*σ_μ_*)) change in the phenotype.

**Figure 3.**
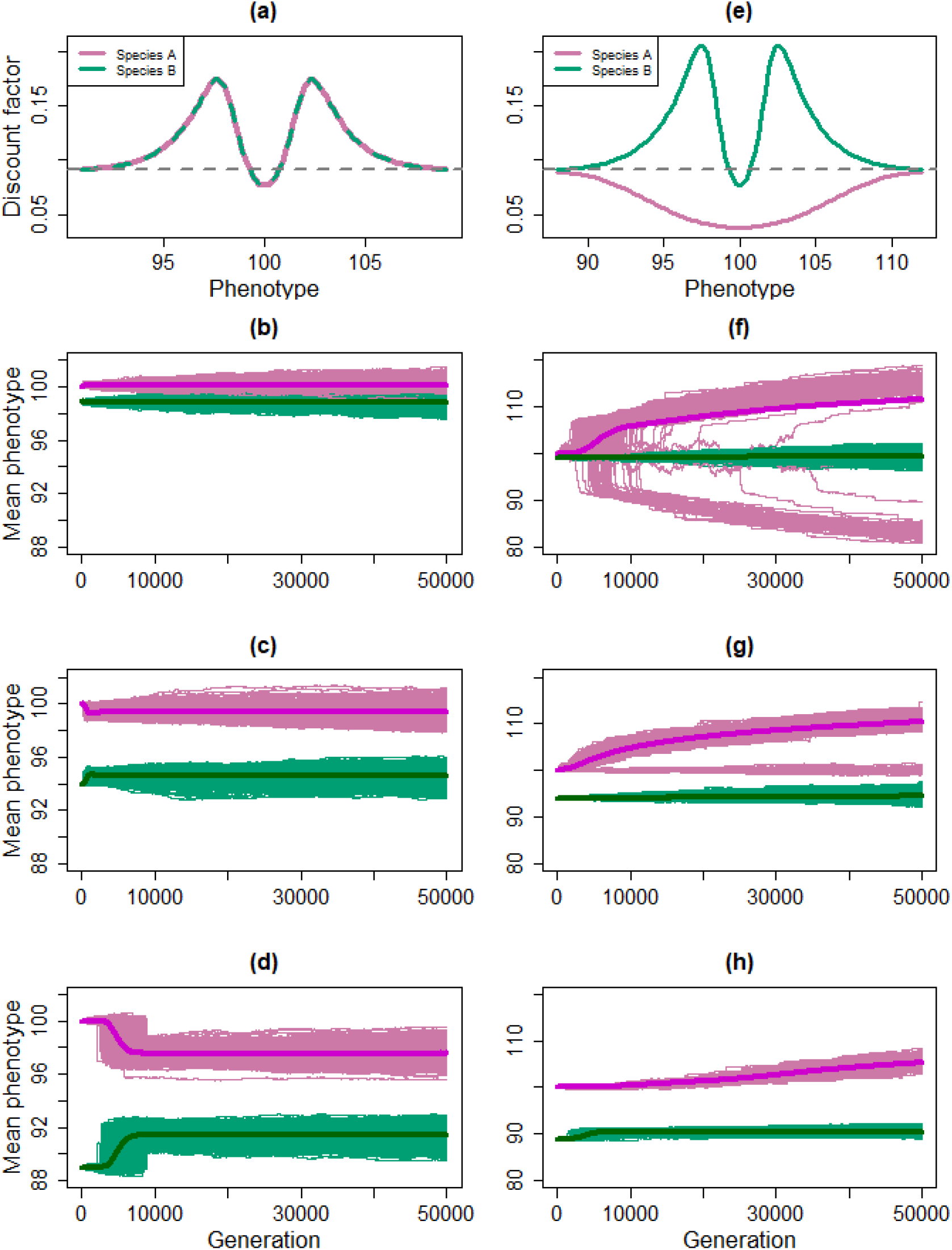
Summary of the evolutionary simulations. First column (a–d) shows scenario 1 where the costs of competition and the benefits of interspecific social information use are symmetric for the two species. Second column (e– h) shows scenario 2 where the costs and benefits are asymmetric for species A and species B so that the information user (species B; green) benefits from using the information, while the information source (species A; magenta) only suffers from interspecific competition with the information user. The first row (a, e) shows discount factors, that is coefficients for converting potential fecundity to realized fecundity in relation to phenotype in the beginning of the simulations (see Section 2, Eq. 2). The grey dashed line represents the level where realized fecundity is independent of interspecific interaction. Rows 2–4 show the trajectories of population mean phenotypes in individual simulations (thin lines) and the mean of all simulations (thick lines) across generations. Simulations of both scenarios were repeated by starting from three different phenotypic distances (cases i-iii in the text). The second row (b, f) shows results of simulations when (i) the initial phenotypes of species A and B were set to 100 and 99, the third row (c, g) when (ii) they were 100 and 94, and the fourth row (d, h) when (iii) they were 100 and 89, respectively. In all simulations presented here b = 1, μ = 0.1, and σμ = 0.1. The other parameter values for the simulations were in scenario 1: F_A_ = F_B_ = 200, a_cA_ = a_cB_ = 1, a_bA_ = a_bB_ = 0.8, c_cA_ = c_cB_ = 1, c_bA_ = c_bB_ = 3, and in scenario 2: F_A_ = F_B_ = 200, a_cA_ = 1.5, a_cB_ = 1, a_bA_ = 0, a_bB_ = 0.8, c_cA_ = 4, c_cB_ = 1, c_bA_ = c_bB_ = 4. In panel g, the simulations where the phenotype of species A remained stable over generations were associated with an immediate extinction of species B and, thus, there was no selection by interspecific interactions affecting species A. Note the different scale of the y-axis between the columns.

We used the model for assessing evolutionary dynamics in two scenarios: (1) the costs of competition and the benefits gained from social information use are symmetric for the two species, and a net cost of competition at high phenotypic similarity changing to a net benefit of information use at sufficient phenotypic difference, and (2) the costs and benefits are asymmetric for species A and species B so that the information user (species B) can benefit from using the information, while the information source (species A) only suffers from interspecific competition with the information user. We repeated simulations of both scenarios by starting from three different initial phenotypic distances between the two species: (i) 1, (ii) 6, and (iii) 11 phenotypic units. In each case (i–iii), we ran 1,000 simulations, each for 50,000 generations, by starting populations of both species from a random number of individuals derived from a uniform probability distribution between 5 and 40 individuals. We also repeated all simulations with three values of *b*, 1, ½ and 1/3, to assess the effect of the strength of joint density dependency on the predictions. To keep the ecological feedback loop via population dynamics comparable among the simulations ran with different values of *b*, we first found a value of *F_i_* that resulted in a population size of approximately 200 individuals when running the model with one species only. The combination of *b* and *F_i_* has a strong effect on population size, so this adjustment was necessary. (Note that the eventual equilibrium population size in single-species simulations depends also on the initial population size, so exact matching of population sizes was not possible in simulations with a random initial population size, but we could avoid population sizes varying with orders of magnitude among simulations ran with different values of *b* with this method.) We always set *F_A_* = *F_B_, μ* = 0.1, and *σ_μ_* = 0.1. We also set the phenotypes of all individuals within a species to be equal (in phenotypic units) in the beginning of the simulations so that *p_•A_* = 100 and *p_•B_* = 99 in case (i), *p_•A_* = 100 and *p_•B_* = 94 in case (ii), and *p_•A_* = 100 and *p_•B_* = 89 in case (iii). All the parameter values are listed in Fig. 3 and the R code for the simulations is provided in Supporting Information (simulation_script.R).

## 3. Results and discussion

### 3.1 Trait divergence

In the first (i) case of scenario 1 (symmetric effects), with the smallest initial phenotypic difference and with all investigated values of *b* (Fig, 4a–c), the model predicts slight character displacement (Fig. 3b) in accordance with competition theory. Trait divergence evolved in less than 5,000 generations and was low in magnitude, because increasing phenotypic difference resulted in a net benefit of interspecific social information use (Fig. 3a), which then stabilized the phenotypic difference between the interacting species (see Supporting Information Fig. S1b for the phenotypic difference). Since social information is expected to be most valuable when provided by information sources that have similar resource needs as the information user (Laland 2004; Seppänen *et al*. 2007; Jaakkonen *et al*. 2015; Szymkowiak *et al*. 2017), the balance between costs of competition and benefits of social information use is sensitive to the phenotypic distance between the two populations (Fig. 3a).

**Figure 4.**
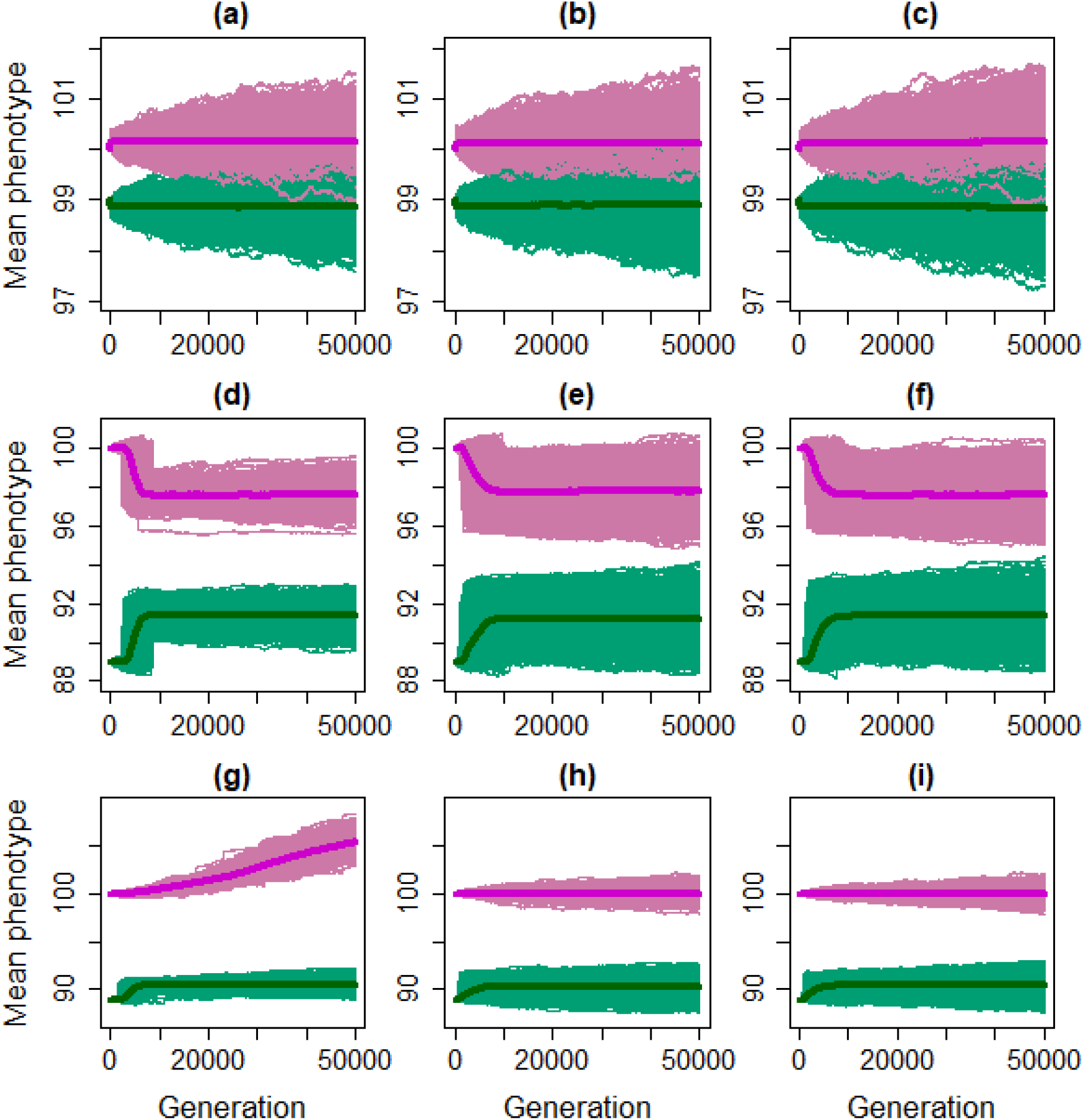
Summary of the effects of parameter b determines the strength of joint density dependency on the three different predicted evolutionary dynamics (divergence, convergence and coevolutionary arms race). The value of b is 1, ½ and 1/3, and the values of F_A_ = F_B_ is 200, 15 and 7 in the first (a, d, g), second (b, e, h) and third (c, f, i) column, respectively. Results of the trait divergence (from scenario 1) are presented in the first row (a, b, c), those of the trait convergence (from scenario 1) in the second row (d, e, f), and those of arms race (from scenario 2) in the third row (g, h, i). Excluding variation in the value of b among columns, the scenario-specific parameter values were the same for each row (1–3) as in Fig. 3b, 3d and 3h, respectively.

Existing literature (Forsman *et al*. 2014; Loukola *et al*. 2020b; Romero-González *et al*. 2020a) provides examples of how SIIU could lead to similar trait divergence as predicted by our simulation model. To acquire social information, individuals need to coexist, which also increases the probability of competition. When ecologically overlapping individuals coexist and compete for the same resources they may actively observe and use the available information produced by putative competitors. This may facilitate the information user to choose a divergent strategy from the information source (Forsman *et al*. 2014; Loukola *et al*. 2020b; Romero-González *et al*. 2020a), leading to a temporary divergence among the individuals in plastic traits (Fig. 2b). Behaviour of unsuccessful individuals should be avoided (reject-the-unsuccessful) to avert possible costly errors increasing the chance to adopt optimal behaviour (Forsman & Seppänen 2011; Romero-González et al. 2020a; but see also Forsman & Kivelä 2021). Rejection behaviour of socially transmitted information from heterospecifics, potentially leading to a temporary trait divergence, has been found in birds (Forsman & Seppänen 2011; Seppänen *et al*. 2011; Loukola *et al*. 2013; Forsman *et al*. 2018; Thorogood *et al*. 2018; Morinay *et al*. 2020) and more recently also in invertebrates (Loukola *et al*. 2020b). Still, there is a lack of evidence for even a temporary trait divergence between the information source and the user in many ecological systems, which may be because of the restrictive conditions under which SIIU can result in a mutual trait divergence (Fig. 3). For SIIU to result in ecological or evolutionary trait divergence, the interacting populations must be initially so close to each other that costs of competition outweigh the benefits of SIIU. If there is a net benefit of SIIU, then divergence cannot continue long even when it occurs (Fig. 3b). Thus, we suggest that SIIU is likely to drive trait divergence only in ecological time scale, and only in plastic traits, if both interacting species share mutual benefits of SIIU. The situation changes if the benefits of SIIU are asymmetric (scenario 2; Fig. 3e), because then stronger – and asymmetric – trait divergence may take place, and the divergence may continue over evolutionary time scale and pertain also to non-plastic traits (Fig. 3f, g).

### 3.2 Trait convergence

In the second (ii) and third (iii) case of initial phenotypic differences in scenario 1 (symmetric effects), the phenotypes of the two species converge with all the tested values of *b* (Fig. 3c, d; Fig. 4d–f). Despite the potential costs of competition, when the phenotypes of the species are initially different enough, both species gain net benefits from social information use and the species’ traits converge if diminishing phenotypic difference results in higher fitness. As expected, increasing initial phenotypic difference increases the time (number of generations) over which convergence continues between the two species (Figs. 3c, d and S1c, d). Contrary to existing competition theories (e.g., Brown & Wilson 1956; Hardin 1960; Costa-Pereira *et al*. 2019; Martin *et al*. 2021; Pastore *et al*. 2021), the recurring beneficial decisions may eventually lead to increased fitness (Farine *et al*. 2015; Keen *et al*. 2020), and therefore, to evolutionary trait convergence between the populations of the two species (Fig. 3c, d). Perfect convergence is not predicted by the model because mutual costs of interspecific competition result in a negative net effect on both species when their phenotypes become too similar (Fig. 3a, c, d). Hence, trait convergence should level off at a point where the net fitness benefit peaks.

As our simulation model predicts, coexistence of individuals of ecologically similar species does not always lead to trait divergence or competitive exclusion. Depending on the intensity of competition, two ecologically similar species may coexist even when their niches overlap in one or more dimensions (Pianka 1974; Martin & Martin 2001; Fox & Vasseur 2008; Chiatante 2019; Martin *et al*. 2021; Pastore *et al*. 2021). Both theoretical (Vellend 2006; Fox & Vasseur 2008; this study) and empirical (Forsman *et al*. 2002; Tobias & Seddon 2009; Seppänen *et al*. 2011; Loukola *et al*. 2013; Tobias *et al*. 2014) evidence show that traits of competing individuals may converge when social information use does not have negative net effects for either of the populations. If putative competitors converge in a specific plastic trait and as a result coexist more than by chance, trait convergence may intensify competition and require differentiation in other plastic traits among the interacting individuals to stabilize similar resource use and coexistence. There is a trade-off between the costs of competition over the same resources and the benefits of social information use (Seppänen *et al*. 2007; Forsman *et al*. 2008; Lee *et al*. 2016) but social information use may allow coexistence of ecologically more similar species than would otherwise be possible (Forsman *et al*. 2002; Loukola *et al*. 2013).

Selective copying of behaviours of successful heterospecific individuals (Laland 2004; Parejo *et al*. 2005; Loukola *et al*. 2013; Laland *et al*. 2020; Romero-González *et al*. 2020a) will result in temporary trait convergence among potentially competing species, similar to our model predictions. SIIU increases reproductive success of the information user in similar environmental conditions as the information source and, given that SIIU does not incur costs for the information source, move the information user towards the information source in the trait space (Fig. 2c). Trait convergence occurs, for example in breeding habitat preference (e.g., Forsman et al., 2002; Parejo et al., 2005; J. T. Seppänen & Forsman, 2007), and may also be a consequence of context-dependent dispersal where breeding habitats are preferred through heterospecific attraction (Mönkkönen *et al*. 1990; Parejo *et al*. 2005; Cayuela *et al*. 2018; Chiatante 2019). With trait convergence, trait similarity should preserve ecological value of the social information for the user since the ecological distance between the information user and the source remains short (Seppänen *et al*. 2007). We propose that trait convergence is the most prevalent evolutionary consequence of SIIU.

### 3.3 Evolutionary arms race

In all cases (i–iii) of scenario 2, with asymmetric costs of competition and benefits of social information use among the two species, species A (information source) is predicted to diverge from species B (information user; Figs. 3f–h and S1f–h) even if the information user would not converge towards the information source. In the case (iii) with the highest initial phenotypic difference of scenario 2, species B first evolves towards species A because it gets fitness benefits from social information use by getting closer to species A in the trait space, but this is followed by species A evolving farther away from species B in response to the increasing costs of competition with species B when *b* = 1 (Fig. 3h). This evolutionary chase suggests potential for evolutionary arms race between species when the costs of competition and the benefits of interspecific social information use are distributed asymmetrically between the species. However, this outcome is strongly affected by the strength of joint density dependency as the phenotypic change between the two species stabilized with *b* = ½ and *b* = 1/3 (Fig. 4g–i), that is, when joint density dependency was weak.

Indeed, the consequences of social information use may be asymmetric between the information source and the information user (Forsman *et al*. 2007; Magrath *et al*. 2015) if one individual suffers negative consequences and the other individual benefits from social information use, or the magnitude of costs and benefits differs between the individuals. For example, there is empirical evidence that the information source bears the costs of the interaction, while the information user gains net benefits from social information use (Forsman *et al*. 2007; reviewed by Magrath *et al*. 2015). This asymmetry in social information use could lead to an evolutionary arms race (Fig. 3h) between the populations of the information source and the user. Arms race occurs if the information source evolves a counter-adaptation to conceal information used by the other species, which, in turn, selects for the evolution of means to overcome those information source counter-adaptations in the information user, followed by the evolution of new counter-adaptations in the information source and so on (Seppänen *et al*. 2007; Loukola *et al*. 2014; Fig. 2d). This way, social information use may enhance the rate of coevolution (Paenke *et al*. 2007; Whitehead *et al*. 2019; Laland *et al*. 2020).

As predicted by our simulation model, interspecific social information use may be one of the mechanisms that has resulted in an evolutionary arms race in interspecific eavesdropping on olfactory cues in bee communication (Nieh *et al*. 2004), alarm call behaviours of multiple species (Blumstein 1999; Keen *et al*. 2020) and passerine breeding habitat selection (Loukola *et al*. 2014). A compelling example of such an arms race appears to occur between competing great tits (*Parus major*) and pied flycatchers (*Ficedula hypoleuca*). Later-breeding flycatchers acquire information on habitat suitability from tit breeding success (i.e., clutch size) and settle near successful tits, plausibly increasing competition (Loukola *et al*. 2013; Forsman *et al*. 2018). As a result, the presence of flycatchers negatively affects the breeding success of great tits (Forsman *et al*. 2007) and, consequently, great tits try to reduce the information parasitism of flycatchers by hiding the information (i.e., covering eggs; Loukola *et al*. 2014, 2020a). Each of the species tries to pursue the most suitable breeding habitat by outrunning the competitor either by gaining or hiding information, possibly leading to coevolution of traits involved in information acquisition and hiding (Loukola *et al*. 2014). To our knowledge, this is the only experimentally demonstrated example of SIIU resulting in a potential arms race between species.

## 4. Community-level consequences of SIIU

SIIU-derived decisions at individual level may generate population-level evolutionary consequences, which, as argued above, may feed back to community dynamics and structure. The prevalence of social information use is dependent on con- and heterospecific densities which affect the intensity of competition (Laland 2004; Seppänen *et al*. 2007; Forsman *et al*. 2008; Jaakkonen *et al*. 2015; Gil *et al*. 2019) and availability of information (Schmidt *et al*. 2015; Kendal *et al*. 2018). Therefore, the decision of the information user (e.g., copy or reject a breeding habitat preference), fitness consequences (i.e., breeding success, survival) for both the information user and the information source, and the balance and continuity of these interactions over generations may affect species’ population sizes leading to species’ aggregations, segregations or in constantly fluctuating community dynamics (cf. Figs 3 and 4). Thus, SIIU may result in higher densities of putative competitors (e.g., heterospecific attraction; Mönkkönen *et al*. 1990) than expected by the competition theory or lower densities of ecologically similar species in their expected habitat (e.g., rejection; Loukola *et al*. 2013). The strength of joint density dependence is also predicted to influence the evolutionary consequences of social information use when the costs of competition and the benefits of social information use are asymmetric between the interacting species (Fig. 4). All the different outcomes of social information use will in turn modify the frequency of potential interactions among species enabling the use of interspecific social information. The community-level effects may also cascade across trophic levels, for example via effects on predation rates. In some cases, SIIU may eventually result in a trait divergence or convergence to the degree that information use no longer pays off, and then we may not find SIIU between the species in question in the present time, although SIIU has been important in the coevolutionary history of those species.

It is likely that potential niche convergence turns into divergence if densities of the interacting species exceed some threshold value (Forsman *et al*. 2008). The balance between the costs of competition and the benefits of social information use is one of the main determinants of the direction of selection (Lee *et al*. 2016). If costs of competition are higher than the benefits of social information use and, consequently, SIIU results in poor fitness of the information user, selection should act against social information use resulting in the evolution of trait divergence among the two species. Although increased competition is one of the main consequences of social information use, even strong negative consequences may not necessarily negate the selection pressure for social information use (cf. Forsman & Kivelä 2021). Furthermore, in a network of interacting species, trait divergence or convergence between two species could lead to divergence or convergence in relation to other species within the same community, altering species’ interactions even further.

SIIU may lead to different, similar or constantly changing adaptations in resource use, through divergence, convergence and arms race, respectively (e.g., in habitat choices, foraging decisions). However, the initial phenotypic distance between the interacting species seems important for the evolutionary outcome; whether the traits of the interacting species start diverging or converging or remain unchanged after the populations come into contact. If SIIU leads to divergence in species’ traits, ecological distance among the species increases, decreasing the value of social information, whereas convergence of traits should increase the value of social information (Seppänen *et al*. 2007). Thus, ecological or evolutionary changes in species’ locations in the trait space – for whatever reason – may facilitate SIIU between new species pairs and break down existing SIIU connections between species. Given the multitude of possible other perturbations to species’ positions in trait space, in dynamic community contexts over evolutionary time, populations encountering conditions triggering SIIU-driven divergence, convergence or arms-race should not be rare. Hence, social information use network among species is expected to be dynamic in both ecological and evolutionary time, highlighting the importance of incorporating these eco-evolutionary dynamics in community ecology.

## 5. Future directions

The theoretical predictions (trait divergence, convergence and arms race) derived from our simulation model should be further tested with experimental studies. Previous studies of social information use have generally concentrated only on the occurrence of copying decisions based on socially transmitted information at the individual level (Danchin *et al*. 2004; Gil *et al*. 2018; Hämäläinen *et al*. 2021b). However, the scope of studies of SIIU should be substantially expanded to better understand the consequences of SIIU at the population- and community-level ecological processes, as well as in the evolutionary time scale. Here we suggest some future avenues.

Expanding empirical research to include new taxa is needed to assess how widespread SIIU is in the animal kingdom. The studies of ecological and evolutionary outcomes of SIIU are still lacking a comprehensive theoretical framework and, thus, extending the ecological and evolutionary theory to include SIIU is called for. Our modelling exercise demonstrates that the effects of SIIU on individual fitness could influence population-level ecological processes, which then translate into selection pressures resulting in a coevolutionary change in the interacting populations. The predictions of SIIU and the current model should be experimentally tested in a broad set of species to assess how widespread SIIU is in the animal kingdom. Ideally, strong inference would require creating experimental designs that control for the learned and genetically determined behaviours in the selective decision-making. Experiments utilizing artificial symbols (Giurfa *et al*. 2001; Seppänen & Forsman 2007) provide one such powerful method where only imagination sets the limits.

Behavioural plasticity supports individuals’ ability to make different selective decision (copy/reject) in different times, based on current and previously observed social information (Mery & Burns 2010; Snell-Rood 2013; Saltz *et al*. 2017). SIIU is also hypothesized to take place consistently through the lifetime of an individual (Forsman *et al*. 2018) similar to intraspecific social information use (e.g., Hämäläinen *et al*. 2021b) and some other behavioural traits such as tendency to explore (Groothuis & Carere 2005) and copying styles (Koolhaas *et al*. 1999). Hence, the success of SIIU should be measured as the net effect gained from social information use over individual’s lifetime. Moreover, it is unknown which traits may serve as information and therefore, also the potential spatial and/or temporal limitations for the information source and the information user. For example, there is a positive association between the size of great tit females and the probability of pied flycatchers to settle near a tit nest over a territory further away (Hämäläinen, R., personal communication). Hence, SIIU with multiple cues of information and time lags are interesting directions for future research.

The consequences and use of social information can be context-dependent (Laland 2004; Kendal *et al*. 2018; Gil *et al*. 2019). Abiotic (e.g., climate and environmental change; Both *et al*. 2004; Samplonius *et al*. 2018) and biotic (e.g., age or sex; Loukola *et al*. 2012) factors may change the context or alter the accuracy of social information (Seppänen *et al*. 2007). Current climate and environmental changes may, for example, lead to asynchrony in nesting times of birds (Both *et al*. 2004; Samplonius *et al*. 2018) or the change in information acquisition strategies of animals (Forsman & Kivelä 2021), and may increase the spatial and temporal distance between species (Seppänen *et al*. 2007), potentially having profound effects on the availability and accuracy of social information. Thus, we suggest identifying the abiotic and biotic contexts that alter the probability and extent of social information use in different circumstances.

Lastly, we encourage to examine the heritability and additive genetic variance in the use of social information. Existing studies show no (Morinay *et al*. 2018) or only weak (Tolvanen *et al*. 2020) evidence for a link between social information use and other heritable social traits, but only few species have been studied in this respect. Still, social information use may be affected by aggressiveness and boldness (Réale *et al*. 2007; Kurvers *et al*. 2010; Morinay *et al*. 2020), and the heritability of these traits (Drent *et al*. 2003; Brown *et al*. 2007; Réale *et al*. 2007) could be reflected in the evolutionary dynamics of SIIU.

## 6. Conclusions

Social information use is a widespread and well-studied phenomenon, yet some key aspects of it remain insufficiently understood. We argue that Selective Interspecific Information Use (SIIU), in which observed behaviours are actively either copied or rejected depending on the perceived success of the source of the information, is a common mechanism of social information use, which has important evolutionary consequences; it may result in trait divergence, convergence or a coevolutionary arms race between interacting species. SIIU may have broader ecological consequences than previously thought, affecting population dynamics and community structure. Maturing the scientific research on SIIU requires connecting SIIU to ecological and evolutionary theory, and such theoretical work is called for.

## Acknowledgements

We thank Mahdi Aminikhah, Tuomas Kankaanpää, Thomas Merckx, Matthew Nielsen and Mahtab Yazdanian for their help in improving previous versions of the manuscript. We also wish to acknowledge CSC – IT Center for Science, Finland, for computational resources. We thank the Unit of Ecology and Genetics at the University of Oulu for funding the artwork in the manuscript. The work of R. Hämäläinen was funded by Societas pro Fauna et Flora Fennica and Finnish Cultural foundations North Ostrobothnia Regional Fund (grant numbers 60182024, 60212359), work of M.H. Kajanus was funded by University of Oulu Kvantum Institute and the Unit of Ecology and Genetics, work of J.T. Forsman was funded by Academy of Finland (projects 122665 and 125720) and Kone Foundation, work of S.M. Kivelä was funded by Academy of Finland (projects 314833, 319898, 345363), work of O.J. Loukola was funded by Academy of Finland (project 309995) and Kone Foundation (project 202010852).

## Glossary

Social information: Observable information resulting directly or indirectly from con- and heterospecific behaviour.
Information source: An individual whose behaviour and/or the consequential fitness outcome can act as a source of information to others.
Information user: An individual who utilizes the information provided by others.
Observed behaviour: Any behaviour of the information source that is observable to the information source.
Selective interspecific information use: The information user selectively copies behaviours or decisions of the information source that result in a high fitness and rejects such behaviours or decisions of the information source that are associated with a low fitness.
Trait divergence/convergence: Divergent or convergent change in species’ trait space, for example as a result of social information use or competition.
Trait space: A multidimensional space whose dimensions are determined by both morphological and behavioural traits.

