## Supplementary Figure S1 for "Ecological and evolutionary consequences of selective interspecific information use"

**Shared first authorship*

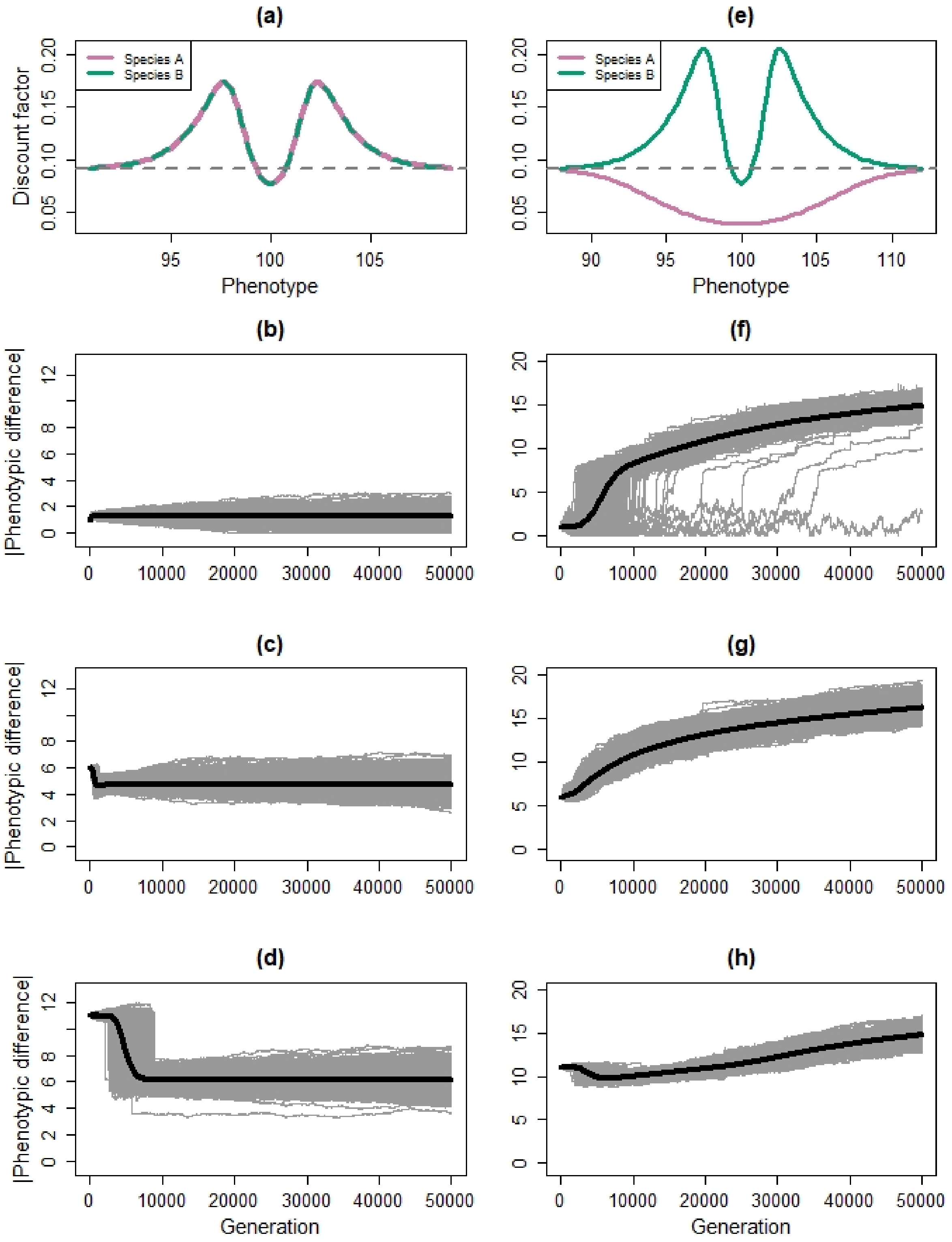


***Figure S1.*** *Summary of the evolutionary simulations. First column (a–d) shows scenario 1 where the costs of competition and the benefits of interspecific social information use are symmetric for the two species. Second column (e– h) shows scenario 2 where the costs and benefits are asymmetric for species A and species B so that the information user (species B; green) benefits*

*from using the information, while the information source (species A; magenta) only suffers from interspecific competition with the information user. The first row (a, e) shows discount factors that is coefficients for converting potential fecundity to realized fecundity in relation to phenotype in the beginning of the simulations (see Section 2, Eq. 2). The grey dashed line represents the level where realized fecundity is independent of interspecific interaction. Rows 2–4 show the trajectories of phenotypic difference between the two populations in individual simulations (solid thin lines) and the mean of all simulations (thick lines) across generations. Simulations of both scenarios were repeated by starting from three different phenotypic distances (cases i-iii in the main text). The second row (b, f) shows results of simulations when (i) the initial phenotypes of species A and B were set to 100 and 99, the third row (c, g) when (ii) they were 100 and 94, and the fourth row (d, h) when (iii) they were 100 and 89, respectively. In all simulations presented here* $b = 1$, $\mu = 0.1$, *and* $\sigma\mu = 0.1$. *The other parameter values for the simulations were in scenario 1:* $F_A = F_B = 200$, $a_{cA} = a_{cB} = 1$, $a_{bA} = a_{bB} = 0.8$, $c_{cA} = c_{cB} = 1$, $c_{bA} = c_{bB} = 3$, *and in scenario 2:* $F_A = F_B = 200$, $a_{cA} = 1.5$, $a_{cB} = 1$, $a_{bA} = 0$, $a_{bB} = 0.8$, $c_{cA} = 4$, $c_{cB} = 1$, $c_{bA} = c_{bB} = 4$. *Note the different scale of the y-axis between the columns.*
